# Population estimate, natural history and conservation of the melanistic *Iguana Iguana* population on Saba, Caribbean Netherlands

**DOI:** 10.1101/2022.05.19.492665

**Authors:** Matthijs P. van den Burg, Hannah Madden, Adolphe O. Debrot

## Abstract

Intraspecific diversity is among the most important biological variables, although still poorly understood for most species. *Iguana iguana* is a Neotropical lizard known from Central and South America, including from numerous Caribbean islands. Despite the presence of native melanistic *I. iguana* populations in the Lesser Antilles, these have received surprisingly little research attention. Here we assessed population size, distribution, degree of melanism, and additional morphological and natural history characteristics for the melanistic iguanas of Saba, Caribbean Netherlands based on a one-month fieldwork visit. Using Distance sampling from a 38-transect dataset we estimate the population size at 8233 ±2205 iguanas. Iguanas mainly occurred on the southern and eastern sides of the island, between 180-390 m (max altitude 530 m), with highest densities both in residential and certain natural areas. Historically, iguanas were relatively more common at higher altitudes, probably due to more extensive forest clearing for agricultural reasons. No relationship was found between the degree of melanism and elevation, and few animals were completely melanistic. Furthermore, we found that body-ratio data collection through photographs is biased and requires physical measuring instead. Although the population size appears larger than previously surmised, the limited nesting sites and extremely low presence of juvenile and hatchling iguanas (2.4%), is similarly worrying as the situation for *I. delicatissima* on neighboring St. Eustatius. The island’s feral cat and large goat population are suspected to impact nest site quality, nest success, and hatchling survival. These aspects require urgent future research to guide necessary conservation management.

## Introduction

The Greater Caribbean region is a biodiversity hotspot known for its high endemism (Mittermeier et al., 1999; Myers et al. 2000; Maunder et al. 2008; Anadon-Irizarry et al. 2012; DRYFLOR 2016), including for reptiles (Kier et al. 2009). At least 7,500 endemic plant species and 880 vertebrate species have been described for this geographically complex region (BEST 2016). Endemic, often geographically restricted, taxa are a major component of biodiversity and a key criterion to conservation valuation and nature management goal-setting. The Lesser Antillean islands of Saba, St. Eustatius and St. Maarten (the Dutch SSS islands) also form part of the Caribbean biodiversity hotspot region. Together these islands support no less than 223 endemic fauna and flora species (32 subspecies, 191 species), of which 35 are endemic to the SSS islands or Saba Bank, 15 are endemic to the adjacent Northern Lesser Antilles, 110 to the whole of the Lesser Antilles and 58 to the combined Lesser and Greater Antilles (Bos et al. 2018). Of the 35 SSS-island endemics, eight are marine, 26 are terrestrial and one is from brackish water (Bos et al. 2018).

Effective conservation requires accurate data on critical variables, e.g. especially considering demography, natural history and population size (Mills 2007). Alarmingly, however, species and population numbers are decreasing globally with such data only being available for a limited number of species, only few of which are reptiles (Meiri and Chapple 2016; Saha et al. 2018). Moreover, over 80% of IUCN-assessed reptiles are categorized based only on range criteria, given the absence of quantitative population data (Bohm et al. 2013; Saha et al. 2018). Conservationists are hence unable to properly evaluate the impact of threats and identify mitigating actions, which is especially troubling for restricted range, numerically-small and genetically-depauperate insular populations.

Endemic species have an elevated extinction risk given they have restricted distributions with small and relatively “closed” populations, and are often ecologically “naive” (Lomolino et al. 2017). Threats include the introduction of new species (rats, cats, goats, raccoons, mongoose, invasive plants), habitat destruction (e.g. coastal development), hunting (e.g. iguanas), illegal trade and stochastic events like hurricanes (Johnson and Winker 2010; Medina et al. 2011; Meléndez-Vazquez et al. 2019; van den Burg et al. 2022a). In the West Indies, 80% of species extinctions have primarily been caused by biological invasions and the region continues to be a “hotspot” of insular extinctions (Leclerc et al. 2018). Since the 16th century the number of non-native vertebrate species has sharply increased (Kemp et al. 2020), which have contributed to the relative high extinction rate of Caribbean reptiles (Slavenko et al. 2016). Currently, 40% of all IUCN-assessed reptiles in the Caribbean region are threatened with extinction (IUCN 2021).

Given the ongoing biodiversity crisis (Ceballos et al. 2017), biologists are attempting to understand global species diversity and intraspecific variation within known species complexes (e.g., Chapple et al. 2008; Ribeiro-Júnior et al. 2022). Indeed, for reptile species, descriptions have exponentially increased during the last two decades (Uetz et al. 2022). In *Iguana* Laurenti 1768, current research aims to understand intraspecific variation within the *Iguana iguana* complex which holds four major mitochondrial clades (Stephen et al. 2013). Although no further genetic data have been published on the extent and boundaries of native populations between these four clades, studies focusing mostly on morphological characteristics have recently proposed two new species, *Iguana melanoderma* and *Iguana insularis* (Breuil et al. 2019, 2020).

Breuil et al. (2020) described *Iguana melanoderma* to include the iguana populations on Saba and Montserrat in the Lesser Antilles, the Virgin Islands, St. Croix, as well as iguana populations in north-eastern continental Venezuela (although no clear boundaries have been described), and adjacent Venezuelan coastal islands (e.g., Isla Margarita, Los Roques and Isla La Banquilla). However, we follow the proposed taxonomy as outlined by the Iguana Taxonomy Working Group (ITWG 2022), in which these populations are considered as part of *Iguana iguana iguana* while awaiting more data from the entire species complex to better assess its taxonomic position.

Irrespective of their taxonomic nomenclature, native populations of keystone species should be protected and studied to better understand their threats and conservation needs. Considering the proposed taxon *Iguana melanoderma* by Breuil et al. (2020), very few data currently exist, presumably given it forms part of the Common Green Iguana complex that is regionally seen as a high-risk invasive species (Knapp et al. 2021). For the Saba iguana population, at least two major threats are known. Firstly, the Lesser Antillean region is known for its high occurrence of nonnative iguana populations, which are spread as cargo shipment stowaways, by hurricanes and during ensuing hurricane-aid campaigns (Censky et al. 1998; van den Burg et al. 2020a, 2021a). As iguanas within the *Iguana* genus, as well as with other Iguanid genera (Moss et al. 2017), can hybridize (Vuillaume et al. 2015), the threat of non-native iguana incursions is evident. Secondly, traders in the exotic global pet-trade are known to have illegally acquired iguanas from Saba which adds considerably to the threat status of this island population (Noseworthy 2017; van den Burg and Weissgold 2020; Mitchell et al. unpubl. data).

In view of the almost total lack of population ecological and conservation information from this native melanistic iguana population on Saba, the objective of this study was to conduct a baseline assessment of the species’ natural history, population size and distribution across the island.

## Materials and Methods

### Study area

This study was performed on Saba, a small Lesser Antillean island (13 km^2^; 17.63°N, −63.23°W; Fig. 1), home to a human population of ~2,000 (Statistics Netherlands 2021). Habitat types were recently mapped by de Freitas et al. (2016) and fall within the Caribbean dry forest biome. The main geological characteristic of the island consists of an 887-m dormant volcano (Roobol and Smith 2004) named Mount Scenery. The volcano causes air uplift and adiabatic cooling, thereby creating a high presence of clouds and mist at elevations above 500-600 m. Below these rainforest-covered elevations, the natural vegetation is drier and less dense, representing xeric Caribbean forest and shrubland. However, large areas of the valleys as well as several hills (e.g., Old Booby hill) are almost devoid of trees due to overgrazing by a large feral goat population, as well as the effects of a disease that wiped out most of the formerly dominant *Tabebuia heterophylla* (de Freitas et al. 2016). Goat abundance and density are especially high in the coastal arid areas on the south and eastern sides of the island, while goat presence is lower in the more forested northwestern region (de Freitas et al. 2016). As permanent beaches are absent on Saba, soil for nesting and incubation of terrestrial breeders (like iguanas and red-billed tropicbirds *Phaethon aethereus*) is primarily available in the arid coastal areas, which are highly affected by the feral goat population, reducing both soil presence and conditions (de Freitas et al. 2016).

**Figure 1.**
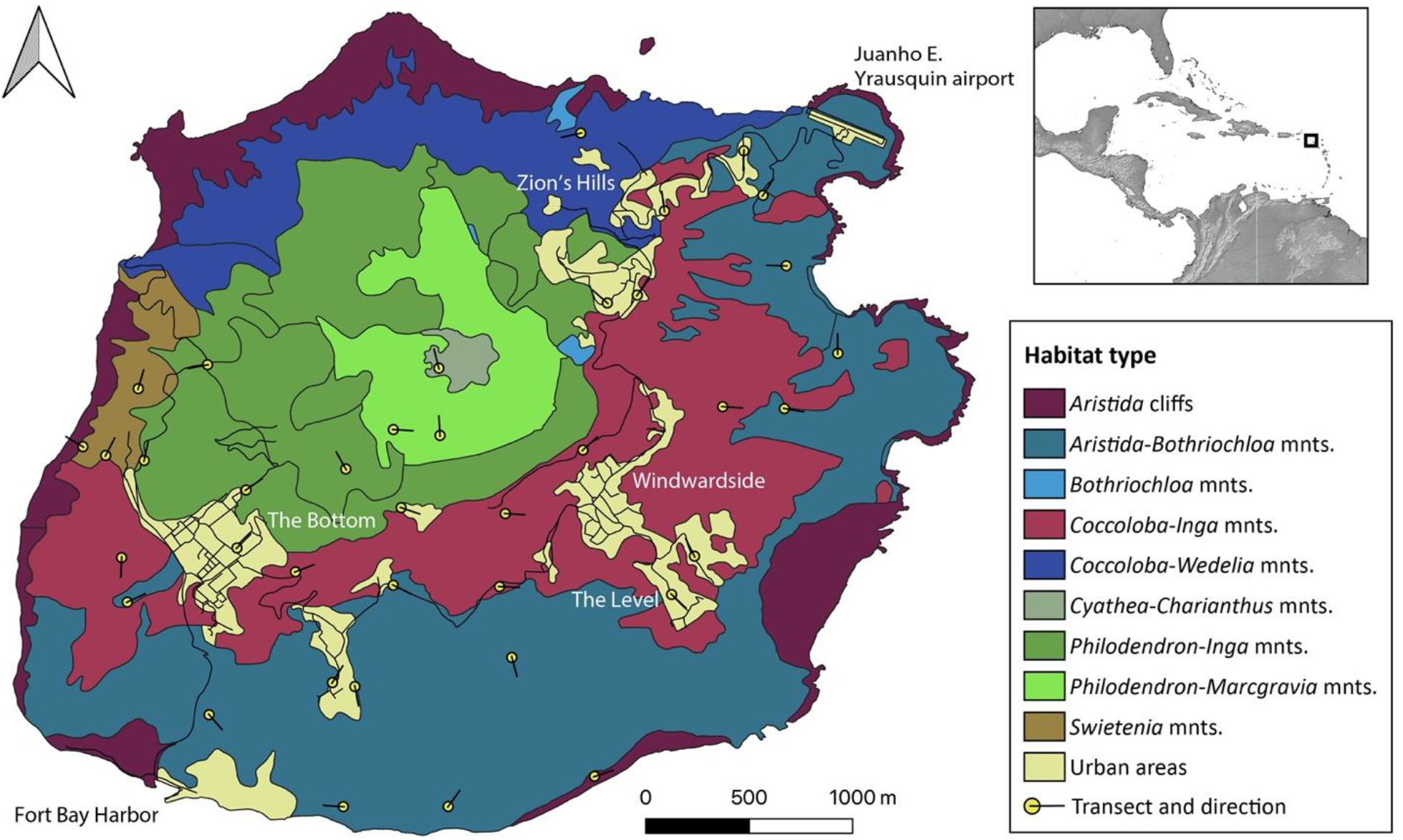
Distribution of transect lines across habitat map of Saba. Names indicate locations on Saba; towns, harbor and airport. Inset map shows location of Saba within Greater Caribbean region. Habitat types follow de Freitas et al. (2016).

### Data collection

In 2021, we collected data during a one-month fieldwork visit to Saba (17 August 2021 – 17 September 2021), using a combination of transect-line surveys and opportunistic sightings. As baseline and trend data on iguana population parameters can be obtained using transect-line surveys (e.g., van den Burg et al. 2022), we implemented this methodology using 100-m transects that were surveyed for iguanas < 25 m on both sides. Transects (n = 38) were distributed over known vegetation types (de Freitas et al. 2016) considering their total surface coverage, while the precise transect locations were assigned randomly within vegetation types prior to the commencement of fieldwork. Each transect was surveyed a minimum of three times, performed between 07.00 – 17.00 h following observations on local diurnal activity patterns. Upon detection, we estimated the perpendicular distance of the iguana from the center of the transect by eye. One vegetation type was not surveyed due to its very limited occurrence (*Bothriochloa* mnts.; de Freitas et al. 2016). The northwestern side of the island was less well presented due to time limitations and limited trail access given goat-eradication efforts.

As Breuil et al. (2020) proposed the importance of two differentiating phenotypic characteristics specifically, but did not provide sample sizes, we collected additional data. First, we took several photographs of all observed iguanas to assess the degree of melanism for adult iguanas, and also the collection of meristic scale counts. Secondly, we measured the maximum height of the tympanum and subtympanic plate using a calliper, to the nearest hundredth of a mm, as well as the snout-vent length to the nearest mm. To collect these data, we opportunistically caught iguanas by hand using a stick and lasso, or using cages.

We further opportunistically recorded iguana presence and GPS locations outside assigned transects. While performing fieldwork we collected data on nest site presence and size, threats (e.g., presence of non-native predators), iguana flight distance, and road kills. Lastly, we informally inquired about local knowledge on the iguana population among local residents and the local park management authority, Saba Conservation Foundation (SCF), in order to obtain additional data and anecdotes about habitat and land use changes over time as well as additional insights.

### Data analyses

We used the vegetation descriptions by de Freitas et al. (2016) to classify vegetation types and to model the habitat-specific abundance of iguanas. We included all vegetation types in which transect surveys were conducted (Fig. 1).

Distance sampling defines the detection function, *g*(*x*), as the probability of detecting an iguana at a distance *x*, whereby for line transects, *x* is the perpendicular distance from the line (Marques et al. 2007). We first examined histograms of the raw data in the R package *Rdistance* (McDonald et al. 2019; R Core Team 2022) to identify clumped detections or outlying observations at minimum or maximum distances in order to apply an appropriate level of truncation. We then applied a right truncation (*w* = 20 m) to the data (Buckland et al. 1993). Following truncation, we analyzed the data using the key function + adjustment terms for the detection function curve, using the Akaike Information Criteria (AIC) to select between competing models (Burnham and Anderson 1998). We also examined the goodness of fit (GOF) of each model. Based on the most parsimonious model, we subsequently derived an estimate of iguana population size in each vegetation type and extrapolated the results across the entire island (13 km^2^). Model precision was determined using 95% confidence intervals and coefficient of variation (CV), with a CV of 20–30% being chosen as the preferred measure of precision. We then used multiple covariate distance sampling to model the detection curve as a function of distance and one or more covariates. The covariates were survey time (minutes), elevation (m), height (of iguana) above ground (m), and weather (0, no clouds; 1, <25% cloud cover; 3, 25–75% cloud cover; 4, 75–100% cloud cover). Low numbers of detections within given surveys made fitting of detection functions difficult and resulted in a low precision of estimates, especially when the data were separated into different vegetation types. Consequently, we pooled distance data across surveys and vegetation types to derive a single estimate of abundance across the entire survey area. Finally, after selecting the most parsimonious model we performed 1,000 iterations to obtain stratified bootstrapped estimates. For the other morphological characteristic, we calculated the ratio between the tympanum and subtympanic plate and assessed its size-dependence using linear regression.

As melanism is the main supposedly distinctive character for *Iguana melanoderma* (Breuil et al. 2020), post-fieldwork we used images of the photographed iguanas to assess the percentage of melanism for each individual. As melanism appears to increase with age (Breuil et al. 2020; this study), we estimated the percentage of melanistic skin surface in larger adults (SVL > 25 cm), distinguishing between head and body. We then classified melanism according to four percentage categories 1) 0–25, 2) 25–50, 3) 50–90, and 90–100%. As iguana photos generally only captured one lateral side, we assumed both sides were identical. Only iguanas with a clearly visible lateral side of the head or/and body were used, others were discarded. Additionally, we recorded whether the lateral spot on the head between the eye and tympanum was melanistic (Breuil et al. 2020).

Data handling and analyses were performed in RStudio Version 1.2.5033 (RStudio Team 2019). Basemaps were created in QGIS 3.8.0-Zanzibar (QGIS.org 2022) and finalized in Adobe Illustrator 25.3.1 (Adobe Inc. 2019).

## Results

### Population census and estimate

Overall, we performed 117 unique surveys, totaling 48 survey hours; respectively, 35 and three transects were surveyed three and four times. During these search efforts we counted 177 iguanas (Table 1), of which 87 were female, 51 were male and 39 were of undetermined sex. Survey-counted iguanas included 168 adults, three subadults, two juveniles and four iguanas of unknown life stage. Among survey transects, most iguanas (n = 44) were encountered on transect #4, located halfway between the main road and coast along the Gilles Quarter trail within *Aristida-Bothriochloa* mountains vegetation. The majority (56%) of transect-counted iguanas were encountered within this habitat type, followed by urban areas (31%) (Table 1).

**Table 1.** Data on survey transects and observed iguanas per vegetation type following de Freitas et al. (2016).

| Vegetation type | Transect count | Iguana sightings |  |
| --- | --- | --- | --- |
|  |  | Transect | Opportunistic |
| *Aristida* cliffs (C) | 2 | 1 | 3 |
| *Aristida-Bothriochloa* mnts. (M7) | 11 | 99 | 172 |
| *Bothriochloa* mnts. (M8) | 0 | 0 | 0 |
| *Coccoloba-Inga* mnts. (M6) | 7 | 14 | 121 |
| *Coccoloba-Wedelia* mnts. (M5) | 1 | 0 | 1 |
| *Cyathea-Charianthus* mnts. (M1) | 1 | 0 | 0 |
| *Philodendron-Inga* mnts. (M3) | 4 | 9 | 9 |
| *Philodendron-Marcgravia* mnts. (M2) | 2 | 0 | 0 |
| *Swietenia* mnts. (M4) | 2 | 0 | 2 |
| Urban areas (U) | 8 | 54 | 172 |
| **Total** | **38** | **177** | **480** |

Opportunistic sightings recorded outside transects totaled 480 iguanas, including 453 adults, 13 subadults, five juveniles, eight hatchlings, and one of unknown life stage (Fig. 2). These concerned 181 female, 138 male and 161 iguanas with unknown sex. All except one hatchling were observed during nocturnal search efforts. In contrast, only a single non-hatchling iguana was observed after sunset. Opportunistic sightings were mostly from urban areas and the *Aristida-Bothriochloa* mnts. vegetation type (each n = 172; Table 1). Considering all observations (n = 657), elevational limits of iguanas ranged from 5–530 m above sea level, with half of all sightings recorded on elevations between 180–390 m. Juvenile and hatchling iguanas represented 2.4% of all iguanas observed on Saba.

**Figure 2.**
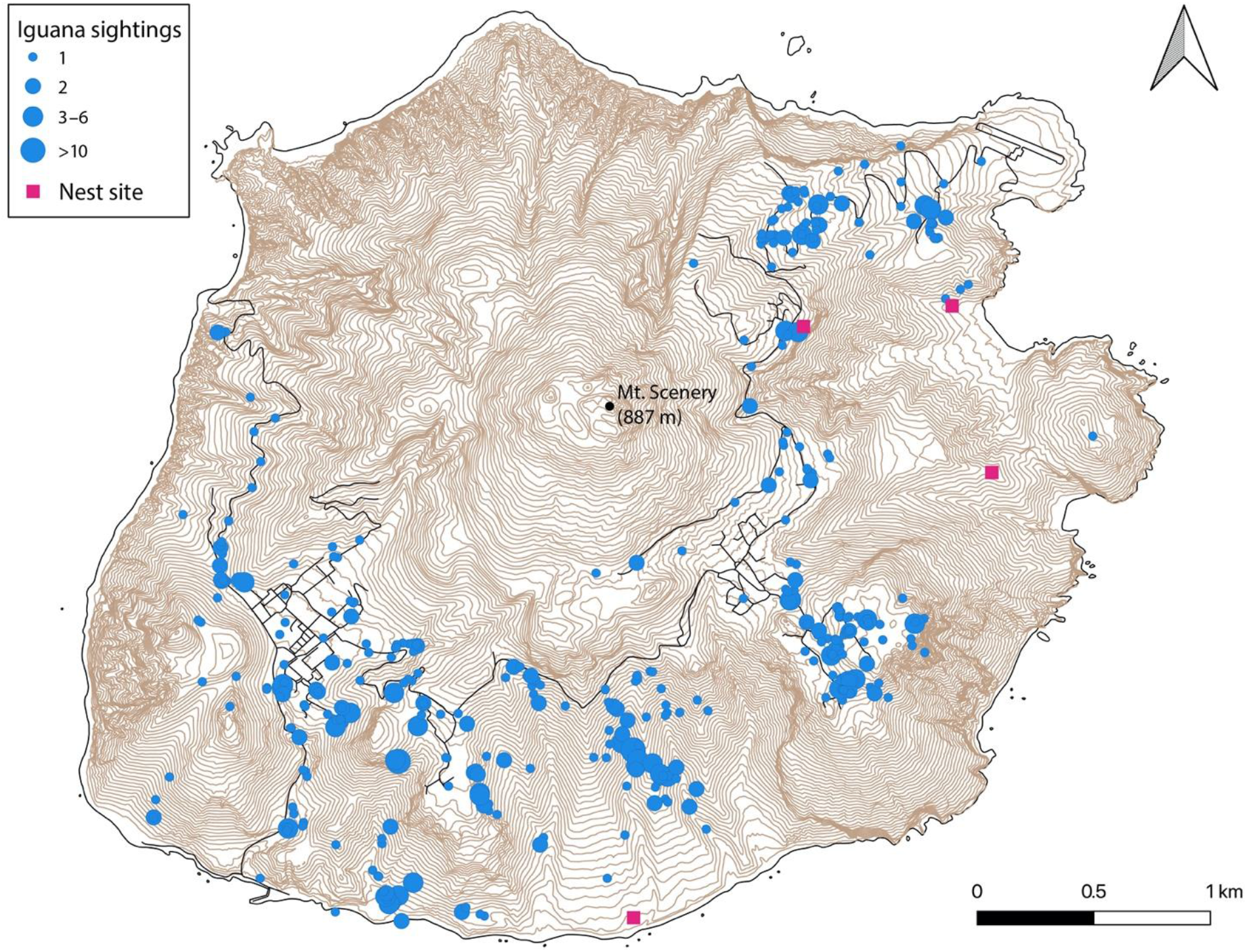
Distribution of sighted iguanas and nest sites across Saba. Data represent both opportunistic sightings and those observed during surveys, n = 657. Contour lines at 10-meter intervals.

Encounter rate varied per vegetation type but was estimated at 23.16 ± SE 5.39 iguanas/transect across the entire survey region. Based on GOF tests and minimization of AIC, the uniform model without series expansion and the half-normal key function with cosine series expansion both provided adequate fit to the distance data (ΔAIC <2; Table 2). Models that included the covariates survey time, elevation + time, and time + weather also received support from the distance data (ΔAIC <2; Table 2). However, we decided to retain the model without covariates (half-normal with cosine series expansion) based on the best GOF and lowest CV. Stratified bootstrapped estimates for this model provided a mean density across the entire survey region of 628.51 ± 168.31 SD iguanas per km^2^ (Table 3). Lastly, bootstrapped abundance estimates suggest a total island iguana population of 8,233 ±2,205 SD animals (adult and subadult).

**Table 2.** Top five-ranked detection models using conventional (CDS) and multiple-covariate (MCDS) distance sampling for iguana line transect surveys on Saba, Caribbean Netherlands, in August - September 2021. Distance data right truncated at *w* = 25 m.

| Method | Key<sup>1</sup> | Series<sup>2</sup> | Covariate<sup>3</sup> | AIC | ΔAIC | $k^4$ | GOF<sup>5</sup> |
| --- | --- | --- | --- | --- | --- | --- | --- |
| CDS | UN | - | - | 523.66 | 0 | 1 | 0.19 |
|  | HN | - |  | 525.49 | 1.83 | 1 | 0.21 |
|  | HN | COS |  | 525.49 | 1.83 | 1 | 0.21 |
|  | HR | - |  | 526.04 | 2.38 | 2 | 0.28 |
|  | HR | SP |  | 526.04 | 2.38 | 2 | 0.28 |
| MCDS | HR | - | Time | 520.67 | 0 | 3 | 0.18 |
|  | HR | - | Elev + Time | 521.18 | 0.51 | 4 | 0.09 |
|  | HN | - | Time | 522.00 | 1.33 | 3 | 0.22 |
|  | HN | - | Time + Weather | 522.04 | 1.37 | 4 | 0.22 |
|  | HR | - | Elev + Time + Weather + Height | 522.72 | 2.05 | 6 | 0.06 |

<sup>1</sup> UN = uniform; HN = half-normal key function; HR = hazard-rate
<sup>2</sup> COS = cosine; SP = simple polynomial series expansion
<sup>3</sup> Time = survey time; Elev = elevation (m), Weather = 0, no clouds; 1, <25% cloud cover; 3, 25–75% cloud cover; 4, 75–100% cloud cover; Height = height of iguana above ground (m).
<sup>4</sup> Number of parameters
<sup>5</sup> Goodness-of-fit

**Table 3.** Stratified bootstrapped estimates of total abundance (left) and density/km^2^ (right) per habitat for the conventional distance sampling model.

| Abundance |  |  |  |  |  | Density (iguanas per km<sup>2</sup>) |  |  |  |  |
| --- | --- | --- | --- | --- | --- | --- | --- | --- | --- | --- |
|  | Mean | SD | LCL | UCL | CV | Mean | SD | LCL | UCL | CV |
| **C** | 2053.16 | 1639.99 | 0 | 5013.24 | 1.00 | 156.73 | 125.19 | 0 | 382.69 | 1.00 |
| **M1** | 0 | 0 | 0 | 0 | 0 | 0 | 0 | 0 | 0 | 0 |
| **M2** | 0 | 0 | 0 | 0 | 0 | 0 | 0 | 0 | 0 | 0 |
| **M3** | 9523.83 | 6862.57 | 0 | 26425.19 | 0.72 | 727.01 | 523.86 | 0 | 2017.19 | 0.72 |
| **M4** | 0 | 0 | 0 | 0 | 0 | 0 | 0 | 0 | 0 | 0 |
| **M5** | 0 | 0 | 0 | 0 | 0 | 0 | 0 | 0 | 0 | 0 |
| **M6** | 6501.53 | 4311.34 | 878.01 | 16434.47 | 0.61 | 496.30 | 329.11 | 67.03 | 1254.54 | 0.61 |
| **M7** | 10592.53 | 3326.75 | 4686.66 | 17826.74 | 0.36 | 808.59 | 253.95 | 357.76 | 1360.82 | 0.36 |
| **Urban** | 19922.09 | 8015.76 | 5442.66 | 36233.81 | 0.36 | 1520.77 | 611.89 | 415.47 | 2765.94 | 0.36 |
| **Total** | 8233.48 | 2204.86 | 4554.74 | 12997.69 | 0.25 | 628.51 | 168.31 | 347.69 | 992.19 | 0.25 |

### Melanism

Overall, we photographed 474 adult iguanas across Saba. Body positioning within the taken photo did not allow us to characterize 139, 162, and 179 individuals, respectively for the lateral spot, head, and body melanism. For head melanism, 79% of characterized iguanas demonstrated over 50% melanism, while for body melanism 81% of characterized iguanas demonstrated less than 50% melanism (Fig. 3). Elevational distribution of the degree of melanism of iguanas showed no significant difference for either head and body melanism (Fig. 3). Visual observation did not indicate any geographic clustering among melanism groups. Only five individuals lacked the facial black spot; these animals were classified as likely non-native given their overall lack of melanism, raised nasal scales, and divergent coloration of body, head and eyes.

**Figure 3.**
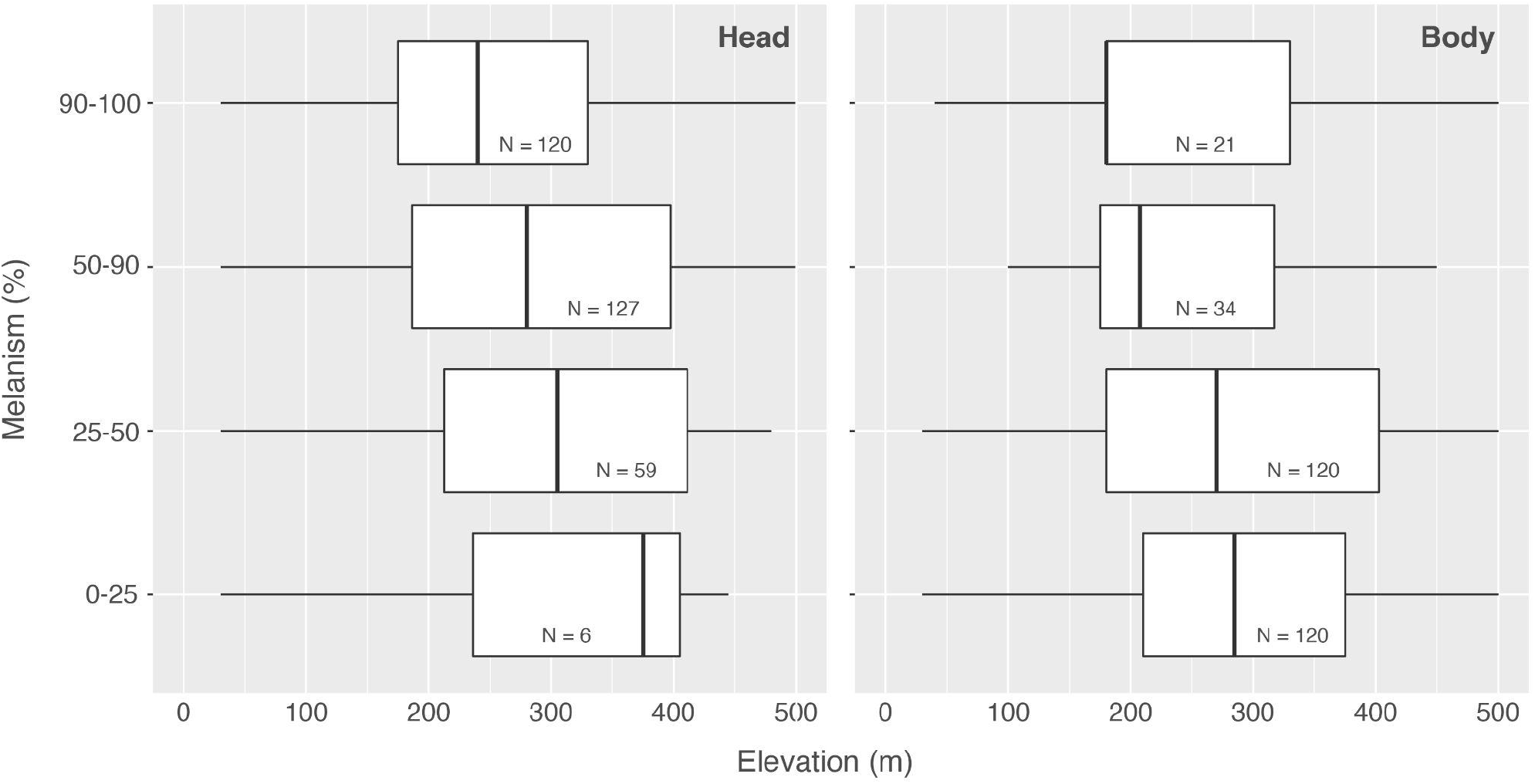
Elevational occurrence of melanism presence in iguanas on Saba, separated by head and body melanism.

### Morphology

We calculated the ratio between the height of the subtympanic plate and the tympanum among 44 adults (>20 cm SVL). This ratio ranged from 1.03–1.84, with two outliers on either extreme (0.92 and 2.22); the ratio was SVL-dependent (Fig. 4). This regression has a *R^2^* = 0.24, and formula y = 0.711 + 0.0021x.

**Figure 4.**
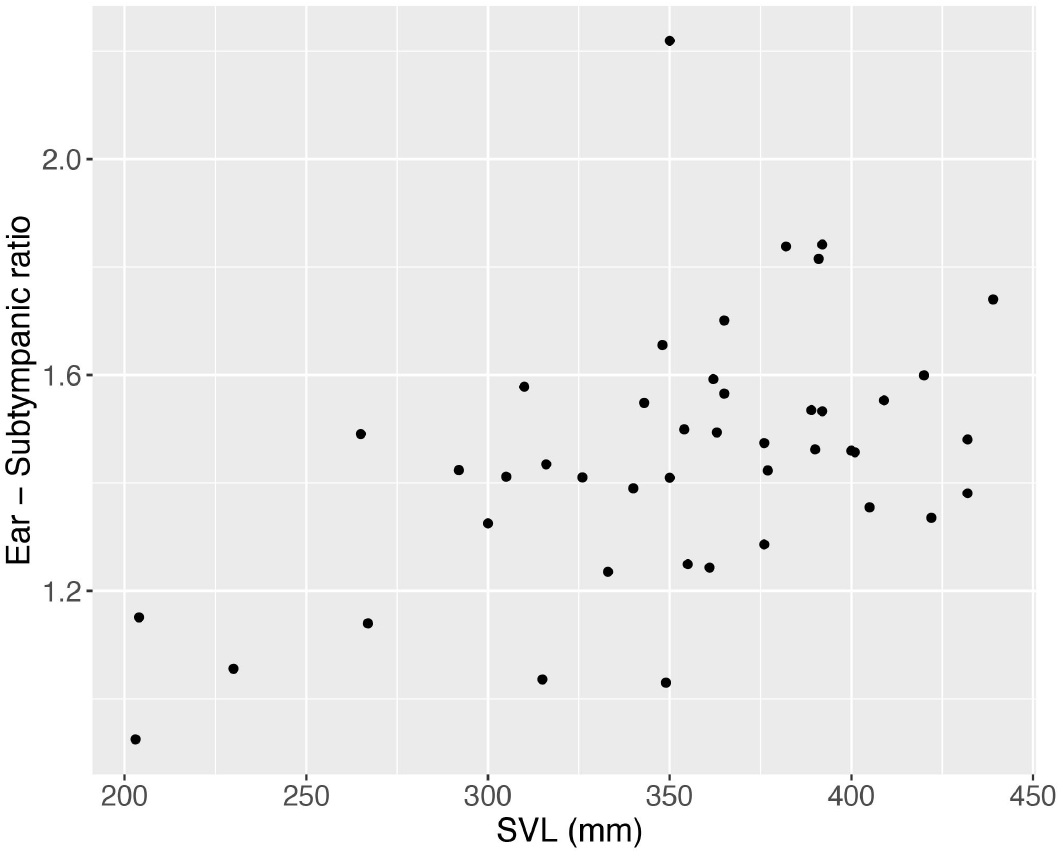
Regression of the ratio between tympanum and subtympanic plate height to snout-vent length for adult iguanas on Saba.

### Other

We actively captured 56 iguanas, as well as collected opportunistic data from two road-killed specimens; 52 adult (>20.0 cm SVL) and 6 hatchling iguanas. Snout-vent length ranged from 9.1–43.9 cm, with female maximum SVL (39.0 cm) being lower than male iguanas (43.9 cm). Emergence holes were encountered at four nest sites, ranging from 2 – 15 holes (Fig. 2), all in areas of shallow soil surface topography (Fig. 2). Flight distance ranged from 0–50 meters.

## Discussion

With only few remaining native *Iguana* populations in the Lesser Antilles, it is important to understand their size, distribution, natural history and threats. Here, we provide the first assessment of the melanistic *Iguana iguana* population of Saba, Caribbean Netherlands. We demonstrate that the population occurs island-wide, except above 530 m, and is estimated at 8,233 ±2,205 SD subadult and adult individuals, results that are in sharp contrast to the preliminary data presented by Breuil et al. (2020). Despite the rather large apparent population size, our assessment suggests that successful recruitment appears to be very low, likely impacted by very limited nest site availability, which is further exacerbated by the large and problematic feral goat population (Lotz et al. 2020). Given the additional stressors of an island-wide feral cat population (Debrot et al. 2014), iguana nesting and recruitment might be severely restricted on Saba. Our work provides a robust baseline for continued research, and highlights essential conservation and future research priorities.

### Distance

Our population estimate for the Saban iguana population is preliminary and based on a limited dataset collected within a short timeframe. While we believe that the true population size may lie in the lower range, especially given the high variability of within-habitat survey estimates and high CV’s (Table 3), we present the first quantitative estimate for this population. Nevertheless, even our lowest estimate far exceeds that of Breuil et al. (2020), who surveyed “some” areas on Saba and Montserrat, resulting in an effective population size estimate of 400 adults for both islands combined. While performing fieldwork on Saba in 2012 the iguana population appeared fairly abundant (A. Debrot, pers. obs.). During the limited time that we conducted surveys, we did not detect iguanas in vegetation types M1, M2, M4 or M5. It is possible that iguanas do not occur in vegetation types M1 or M2, whose high altitudes (> 550 m) and lower temperatures may not be favored. However, we are of the opinion that dense vegetation in these areas makes detection more difficult than in habitats at lower elevations, and we recommend that any future comparative assessments do not exclude these areas. No iguanas were detected in vegetation type M4 during fieldwork, however they are considered to occur here in low densities based on opportunistic sightings (van den Burg pers. obs.). Similarly, no iguanas were detected in M5, however access to the area was severely restricted due to ongoing goat removal actions. Iguanas were detected in vegetation types C, M3, M6, M7 as well as urban areas, suggesting they can persist in fragmented and degraded habitats and among humans, despite the associated risk of mortality that this may bring e.g., vehicles, fences, cats and dogs (Smith et al. 2007; Knapp et al. 2016; van den Burg et al. 2018a).

This long overdue population estimate serves as an important preliminary baseline for longterm monitoring. For future comparison it is important to note that data collection occurred four years after the 2017 hurricane season, which was the strongest-recorded Atlantic hurricane season on record and strongly affected iguana populations on other islands (Schultz et al. 2018; van den Burg et al. 2022a). Similarly, on Saba, both forests and the Red-bellied Racer population were affected by hurricanes during 2017 (Eppinga and Cucko 2018; Madden and Mielke 2021). Therefore, our data could represent those of a recovering population.

### Distribution

The iguana population is present throughout the island of Saba, to a maximum altitude of around 550 m (Figs. 2–4). The majority of the population occurs between elevations of 180–390 m, in habitat types M6, M7, and in urban areas (de Freitas et al. 2016). Although we were unable to visit the northwestern region as mentioned above, interviews with on-island organization SCF and numerous residents indicated that iguanas do occur in that area; though likely at low densities given this represents the leeward side of Mt. Scenery. We thus argue that according to IUCN mapping guidelines (IUCN 2021), the species is present throughout the island, except at altitudes higher than 550 m, and that the initial distribution as presented by Breuil et al. (2020) in figure 12B, is significantly lower than our assessment but was based on limited data.

Lazell (1973) indicated that iguanas were common across the island, including at high elevations around the summit of Mt. Scenery (887 m). This no longer is the case. While Breuil et al. (2020) observed iguanas at a maximum elevation of 500 m at The Level, the authors mentioned that at similar heights (400–500 m) on Mt. Scenery a population of iguanas could not be present due to high moisture and cloud presence. Indeed, the climate appears to differ at similar heights on The Level and Mt. Scenery, due to a difference in wind and sun exposure and forest cover. Further, we lack photographic material from the 1960s during which Lazell visited Saba, which would allow some insight into the presence of degraded, heterogeneous habitats that could provide more opportunities for basking and thereby promote iguana presence at higher elevations. The ability for iguanas to bask is essential to allow proper digestion of their vegetarian food by intestinal flora (van Marken Lichtenbelt 1992). However, data from the 19th century indicates that the largest extent of the Dinzey plantation did reach towards the summit along the western slope, seemingly to ~800 m (Fig. 4 in Espersen 2017). De Palm (1985) also indicates that up until the early 20th century most of the 200 ha of land suitable for small-scale agriculture was located at higher altitudes but by the 1980s only 64 ha of this land was still in use. Hence, it seems likely that at the time of Lazell’s visit, habitats on the west side of the summit were still considerably less-vegetated with secondary forest cover than today, which may have favored iguana presence at higher altitudes.

### Melanism

As nomically evident, Breuil et al. (2020) described *I. melanoderma* with melanism as the most distinct characteristic. However, the extent of melanism per individual differs and increases with age, though a black spot between the eye and tympanum is always present (Breuil et al. 2020). Here we provide additional data on melanism in the Saba population, which is regarded as more melanistic than the population on Montserrat. These data demonstrate that melanism is most clearly represented on the head, with 79% of all photographed and characterized adults for this feature (n=312) showing >50% melanism. Contrarily, melanism is much more limited on the body as 81% of all photographed and characterized adults for this feature (n=295) showed less than 50% melanism. Although numerous adults appear to have body melanism, often the scales are actually dark brown, but might also be dark reddish.

The characteristic black facial area as assigned to *I. melanoderma* by Breuil et al. (2020) was absent in five individuals on Saba, all of which also showed a divergent head and body coloration. The genetic origin and species status of four of these iguanas is currently being assessed using genetic methodology (van den Burg et al. 2021b), and will be discussed elsewhere.

Using open-source images on the internet, Breuil et al. (2020) stated that it is “clear” that non-native iguanas are present on Isla Margarita (Venezuela). However, we have demonstrated here that not all iguanas meeting the “black spot” criteria for *I. melanoderma* are clearly melanistic, and considerable variation is evidently present. Hence, given this seemingly inherent variability without any genetic data, we suggest that confirming the presence of non-native iguanas based on a limited number of open-source photos is tenuous at best. Alternatively, iguana populations in the geographical source region of the Saba and Montserrat populations (insular and continental Venezuela) may be inherently phenotypically variable in this geographically complex region, with periods of isolation and connectivity due to sea level changes during the ice-ages (Mayle et al. 2009).

Melanism can provide an adaptive advantage to ectotherms in cooler climates, as well as provide UV protection at high elevations (Clusella Trullas et al. 2007; Reguera et al. 2014). Our data, however, do not support such explanatory mechanisms for these melanistic populations given the absence of a trend between melanism % and elevation on Saba (Fig. 3). We thus propose that the most parsimonious reason these iguanas exhibit melanism is that the population originates from melanistic iguanas from the northern region (mainland or islands) of Venezuela, as suggested by genetic data (Stephen et al. 2013).

### Morphology

Breuil et al. (2020) state that the ratio between the height of the subtympanic plate to the tympanum in *I. melanoderma* reaches 2–2.5 in large adults. However, the authors provide no mention of sample size or measure methodology, nor is the definition of a ‘large adult’ provided. Here, we assessed this ratio among 44 adults (>20 cm SVL) and found that only a single male (35 cm SVL) had a ratio higher than 1.84; 2.22 (Fig. 4). Considering the retrieved regression between the ratio and SVL (see Results), an iguana would have a ‘subtympanic plate:tympanum’ ratio of two for a SVL of 614 mm. Although Breuil et al. (2020) might have captured larger iguanas as present in our dataset, maximum SVL sizes for the *Iguana iguana* complex are ~45–58 cm SVL (Fitch and Henderson 1977; Dugan 1982; McCranie et al. 2005), and therefore a ratio of 2–2.5 appears unrealistic. Breuil (2013) proposed a set of morphological characters useful to identify between *I*. *delicatissima* and *I*. *iguana*, their hybrids, and several island populations of the later species complex; these were proposed as “determinable from photographs” among which the ‘subtympanic plate:tympanum’ ratio. Although photographs have been used for morphological purposes by other authors (Stevens et al. 2008; Lehtinen et al. 2020) as well as for distribution studies in *Iguana iguana* (van den Burg et al. 2020b; Mo and Mo 2022), we question their use for non-meristic and non-color characters. Even if photographs were taken perfectly perpendicular to the head of the iguana, the tympanum and subtympanic plate would not be in the same 2D-field relative to the camera lens. When doing morphometric comparisons using 2D photographs, a key assumption is that all measurements lie in the same plane relative to the camera. However, especially large (dominant) adults have swollen jaws which ‘push’ the subtympanic plate outwards towards the camera, effectively exaggerating its size compared to the tympanum.

### Clutch size

Clutch size within *Iguana*, as for most reptilians, is SVL dependent but differs per population seemingly due to climate and food availability (Novosolov et al. 2013). For example, iguanas in Central America lay much larger clutches than iguanas on the arid ABC islands (van Marken Lichtenbelt and Albers 1993), and *I. delicatissima* x *I. iguana* hybrids lay larger clutches than pure *I. delicatissima* (van Wagensveld and van den Burg 2018). For the melanistic populations on Saba and Montserrat, only Blankenship (1990) provides some information (clutch size 15–30 eggs) but without an indication of sample size. We dissected a single road-killed gravid female (32.5 cm SVL) found at Lower Zion’s Hill that had 29 well-developed eggs. As the maximum size of females is larger, maximum clutch size within this species is expected to be > 30.

### Nest sites

Successful nesting and recruitment are essential for population size maintenance and long-term survival (Bock et al. 2016; Warret Rodrigues et al. 2021). Although this study was not specifically aimed at identifying communal nest sites, we did survey for and record sites with hatchling emerging holes, four of which were encountered. Therefore, these sites appear to be limited on this volcanic island, as is the case for *I. delicatissima* on neighboring St. Eustatius (Debrot et al. 2013). Both islands are occupied by large feral goat populations that are known to aggregate on and destroy iguana nesting sites (Diaz 1984; Alberts 2004). As all communal nesting sites were located only in sparse areas of shallow-sloping surface topography (Fig. 2), these data provide useful insights for high-priority follow-up studies as to which areas may harbor additional nesting sites.

### Conservation

The Convention on Biological Diversity (CBD) requires member states to identify, describe and monitor important components of biodiversity, including ecosystems and diverse or special habitats as well as threatened and endemic species. The latter represent a unique contribution of any particular region to global biodiversity and are often abundant on or around islands due to genetic isolation which allows them to develop unique traits. Not surprisingly, islands play an important role in generating (as “cradle”) and conserving (as “museum”) biodiversity worldwide (Gascuel et al. 2016), but island endemics are typically more vulnerable to extinction (e.g., Biber 2002; Kouvari and van der Geer 2018; Leclerc et al. 2018).

The IUCN assesses the conservation status of plant and animal species worldwide (IUCN, 2021). However, most rare and endangered island endemics are not included in the assessments due to data deficiency or a perceived low priority (Leclerc et al. 2018). Not surprisingly, of the 223 endemic (sub)species on the SSS islands, assessments are only available for 42 (Debrot et al. 2018). Therefore, basic population assessments that allow IUCN evaluation regarding the status of these species are urgently needed to provide important context for such conservation priority-setting.

### Feral cats

Feral cat populations form a threat to many species (Woinarski et al. 2015; Doherty et al. 2016), including Iguanidae species, as hatchling and juvenile iguanas are especially vulnerable to cat predation (Iverson 1978; Mitchell et al. 2002; Wilson et al. 2004; van den Burg et al. 2018a). Debrot et al. (2014) studied the effect of cats on the Saban Red-billed Tropicbird population using a combination of surveys, camera trapping and faeces analyses. Cats were confirmed to be present island-wide, with higher densities at lower dry elevations, especially close to the landfill area east of the Fort Bay harbor. Although these authors encountered iguana remains in 8.5% of 94 analyzed cat scat samples, no study has yet focused on the population-wide impact of cat presence on the Saba iguana. As cats are numerically more abundant at the lower levels of the island where the vegetation is more amenable for effective hunting by this terrestrial predator (McGregor et al. 2015), it remains to be seen what role cats may play in the lower iguana densities documented for the lower altitudes of the island. Alarmingly, as nesting sites are also predominantly expected at low elevation, given temperature and soil characteristics, higher presence of cats at these elevations could explain the extremely low proportion of juvenile and hatchling iguanas observed during our study period. Although some data on hatchling ecology, movement and survival is present from mainland *I. iguana* populations (e.g., Bock 1984), such data remain lacking for insular islands but would be valuable for conservation management purposes. Finally, a cat-removal campaign that followed Debrot et al. (2014) was only temporary and local in focus, suggesting that feral cats still occur across the island, however recent data are lacking and should be the focus of a future research effort.

### Development

Coastal development threatens iguana populations throughout the Lesser Antilles (van den Burg et al. 2018b; Breuil et al. 2019). Not only are the warmer, dryer and less-vegetated coastal zones generally better habitats for the iguana in terms of sunning opportunities but on many islands the coastal zone has better soil accumulations for nesting. On several islands, the females of this species are known to migrate long distances (e.g. 2 km) to the coast to lay eggs (Gatun Lake islands of Panama, Dominica, Chancel island in Martinique and îles de la Petite Terre) (Bock and McCracken 1988; Breuil 2002; Knapp et al. 2016). On such islands, suitable habitat for laying appears to be limited by both vegetation and geology, and coastal beaches are prime nesting habitat.

On Saba, well-developed plans exist for the construction of a new harbor, located at Black Rock on the central-southern side of the island. This area is highly heterogeneous, with some sites having only a grass-overgrown substrate, while others have larger boulders as well as trees where we observed numerous iguanas (Fig. 2). As reported, an environmental impact assessment mostly focussed on coral and the Red-billed Tropicbird population is insufficient, as long as a robust iguana assessment for this area remains pending.

### Non-native iguanas

Non-native *I. iguana* are widespread in the Greater Caribbean region, especially in the Lesser Antilles (Knapp et al. 2021). The presence of several non-native iguanas on Saba is alarming but not surprising given the extensive occurrence of this species on St. Maarten (Powell et al. 2011), which has also reached neighboring St. Eustatius (van den Burg et al. 2018c; Debrot et al. 2022). As we await CITES documentation for sample shipment and genetic analyses of both samples from Saba and Montserrat, these will be discussed elsewhere (van den Burg et al. unpublished data). Clearly, however, improved biosecurity is necessary to prevent further incursions, especially considering the projected increase in shipments for the harbor construction (Hulme 2009), as well as the tendency to reduce biosecurity measures during aid campaigns in the aftermath of natural disasters (van den Burg et al. 2021b).

### Pet trade

Patterns within the exotic reptile pet-trade are largely driven by species rarity (Janssen 2021). Given the uncommon morphology of iguanas from Saba and Montserrat, there is high interest from the illegal reptile market, and melanistic iguanas are currently sold around the globe (Noseworthy 2017; van den Burg and Weissgold 2020). Recent genetic analyses of some captive melanistic iguanas confirms their origin from Saba or Montserrat (Mitchell et al. unpubl. data). However, no CITES export permit of live iguanas from Saba and Montserrat exists (CITES 2022). Indeed, flaws within the permitting system do not prevent all illegal activity, and some traders mask illegal trade through statements of captive breeding (Janssen and Chng 2018; van den Burg and Weissgold 2020), which is why a trading ban on live *Iguana* sp. from the Caribbean was recently proposed (van den Burg et al. 2022b). Given the predicted increase in cargo traffic from St. Maarten once harbor construction commences, as well as following its completion, we recommend that all harbor, border and customs officials on both islands receive additional training and support in order to serve as a first line of defense against further incursions on Saba.

In contrast to most Lesser Antillean islands, several of which were connected and once harbored various now-extinct lineages of large terrestrial vertebrate species (McFarlane et al. 2014; Brace et al. 2015), Saba is a relatively young volcano that has never been connected to any other islands in the geological past (Roobol and Smith 2004). As such, the melanistic iguana is the only large terrestrial vertebrate ever believed to have lived on the island, which accords it a unique position within the island’s ecology. Understanding critical population variables is hence essential to identify the conservation needs and effectively manage this keystone species. The baseline data provided here form a crucial starting point for long-term monitoring given the ongoing threats from feral mammals, non-native iguanas and increasingly intense hurricane seasons.

## Acknowledgments

This manuscript is part of the Wageningen University BO research program (BO-43-117-006) and was financed by the Ministry of Economic Affairs, Agriculture and Innovation (EL&I) under project number 4318100346-1. We thank the Saba Conservation Foundation for on-island logistical support throughout our project, and the Executive Council for permitting this research (663/2021).

## Conflicts of Interest

The authors declare no conflicts of interest.

## Notes

### Competing Interest Statement

The authors have declared no competing interest.

